# Cardiac interoception is enhanced in blind individuals

**DOI:** 10.1101/2022.05.02.490293

**Authors:** Dominika Radziun, Maksymilian Korczyk, Laura Crucianelli, Marcin Szwed, H. Henrik Ehrsson

**Author notes:** (DR); (MK); (LC); (MS); (HHE).

## Abstract

Blind individuals have superior abilities to perform perceptual tasks that rely on exteroceptive information, since visual deprivation is associated with massive cross-modal plasticity. However, it is unknown whether neuroplasticity after visual loss also affects interoception, i.e., the sensations arising from one’s inner organs that convey information about the physiological state of the body. Herein, we examine the influence of blindness on cardiac interoception, which is an interoceptive submodality that has important links to emotional processing and bodily self-awareness. We tested 36 blind and 36 age-and sex-matched sighted volunteers and examined their cardiac interoceptive ability using a well-established heartbeat counting task. The results showed that blind individuals had significantly higher accuracy in perceiving their heartbeat than did individuals in a matched sighted control group. In contrast, there were no significant differences between the groups in the metacognitive dimensions of cardiac interception or the purely physiological measurement of heart rate, thereby underscoring that the improved accuracy likely reflects a superior perceptual sensitivity to cardiac interoceptive signals in blind individuals. We conclude that visual deprivation leads to enhanced interoception, which has important implications for the study of the extent of massive cross-modal plasticity after visual loss, understanding emotional processing in blind individuals, and learning how bodily self-awareness can develop and be sustained in the absence of visual experience.

## Introduction

Lack and loss of vision are associated with massive cross-modal plasticity (see Frasnelli, Collignon, Voss, & Lepore, 2011). Neuroplasticity, which occurs after sensory deprivation, can lead to enhancements within one or more senses to compensate for the lack of another sense (see Merabet & Pascual-Leone, 2010; Renier, De Volder, & Rauschecker, 2014; Singh, Phillips, Merabet, & Sinha, 2018). In line with this, numerous studies have found that blind individuals show superior performance on perceptual tasks that involve processing exteroceptive information, i.e., stimuli originating outside of the body. Within the auditory modality, blind individuals have been found to have enhanced abilities in spatial hearing both in near (Lessard, Paré, Lepore, & Lassonde, 1998; Röder et al., 1999) and far space (Voss et al. 2004; Battal, Occelli, Bertonati, Falagiarda, & Collignon, 2020), as well as superior pitch discrimination (Gougoux et al., 2004). In the case of the tactile modality, blind individuals have been shown to have enhanced acuity (Goldreich & Kanics, 2006; Wan, Wood, Reutens, & Wilson, 2010), as well as superior tactile symmetry perception (Bauer et al. 2015). Finally, in the olfactory modality, blind individuals have been found to have a lower odor detection threshold (Cuevas et al., 2010; Beaulieu-Lefebvre, Schneider, Kupers, & Ptito, 2011; but see also Sorokowska, 2016). All these sensory enhancements facilitate blind people’s interactions with “the outside”, i.e., the external environment. However, interoception, i.e., sensing oneself from “the inside”, which is crucially important for maintaining bodily awareness and emotional processing, has not yet been investigated in blind individuals.

Interoception, in its classic definition, is the sense of the internal state of the body, which originates from the visceral organs (see Sherrington, 1948). More recent accounts frame interoceptive signals more broadly, including stimuli transmitted through lamina I of the spinal cord, e.g., sharp and burning pain, innocuous warmth and cold, itch, or affective touch, which is information that helps the organism maintain an optimal internal state through homeostatic mechanisms (see Purves et al., 2019; see also Craig, 2002; Björnsdotter, Morrison, & Olausson, 2010; Fotopoulou & Tsakiris, 2017; Crucianelli & Ehrsson, 2022). Among the interoceptive submodalities, the heartbeat is one of the most studied signals (see Khalsa et al., 2018). Cardiac interoception is believed to play an important role in bodily awareness (Herbert & Pollatos, 2012) and emotional functioning (Critchley & Garfinkel, 2017). Alterations in this interoceptive submodality have been described in autism (Garfinkel et al., 2016a) and schizophrenia (Ardizzi et al., 2016).

This experiment aims to investigate the potential influence of blindness on cardiac interoception. To quantify the objective ability to perceive heartbeats, we used the classical heartbeat counting task (Schandry, 1981). Furthermore, to gain a richer understanding of cardiac interoception both at the perceptual and metacognitive levels, the present article follows the dimensional model of interoception introduced by Garfinkel and colleagues (2015; see Suksasilp and Garfinkel [2022] for the revision of the model). This model distinguishes three major dimensions of interoception. The first is interoceptive accuracy, which is an objective performance on a behavioral test consisting of monitoring one’s own physiological events. In this paper, this concept refers to the accuracy in the heartbeat counting task (Schandry, 1981), in which individuals count their heartbeats for a given amount of time. The second is interoceptive sensibility, which is the participant’s assessment of their own interoceptive experiences as obtained by self-report. In this paper, this concept is defined as the result of the Multidimensional Assessment of Interoceptive Awareness [Mehling et al., 2012] questionnaire, which is a measure relating to a spectrum of internal bodily sensations. The third is interoceptive awareness, which is the degree to which interoceptive accuracy correlates with confidence in task response. In this paper, this concept is defined as the correlation between heartbeat counting task performance and the confidence ratings obtained after every trial of the task. Additionally, to examine another dimension of participants’ reflection on their abilities, we obtained the participants’ beliefs about their performance both before and after completing the task. Some interoceptive dimensions have been found to correspond, and others to dissociate, with the dissociations being especially prevalent in clinical populations (e.g., Garfinkel et al., 2016a; Palser, Fotopoulou, Pellicano, & Kilner, 2018, 2020; Jakubczyk et al., 2019; Rae, Larsson, Garfinkel, & Critchley, 2019). Therefore, investigating all the dimensions of interoception instead of one (for example, accuracy only) is important for discussing potential clinical implications of the study.

Given the existence of reports showing the involvement of somatosensory mechanoreceptors in cardioception (Macefield, 2003; Knapp-Kline, Ring, Emmerich, & Brener, 2021), we also included a control task, namely, the grating orientation task, which is a well-established measure of tactile acuity (Johnson & Phillips, 1981). By including this task, we could assess to what extent the potential difference in the ability to detect heartbeats is specific to cardiac interoceptive accuracy itself and not due to the influence of superior tactile acuity of blind participants (e.g., Goldreich & Kanics, 2003; Alary et al., 2009).

Our study is, we believe, the first to look at the relationship between blindness and cardiac interoception, as well as visceral interoception in general. Our hypothesis was that cardiac interoception is enhanced in blind individuals and, thus, that blind individuals would perform better than sighted individuals in the heartbeat counting task. We did not have specific predictions regarding the remaining interoceptive dimensions, as these were included for exploratory purposes. The overarching goal of this study was to take the first step toward understanding how the absence of vision influences interoception, which could have important implications for advancing our understanding of the role of visual experience in bodily self-awareness and emotional processing.

## Methods

### Participants

Thirty-six blind and 36 sighted individuals (age range: 22-45 years, mean age: 33.42 years in the blind group, 33.19 in the sighted group; 19 males and 17 females per group) participated in the study. A sighted, sex- and age-matched participant was recruited for each blind participant. Both blind and sighted participants were invited to take part in the study through multiple recruitment channels to make the samples representative and to balance any potential bias one channel might introduce. All subjects reported that they had no additional sensory or motor disabilities. The exclusion criteria included having a history of neurological or psychiatric disorders.

For all blind participants, blindness was attributed to peripheral origin and was not associated with other sensory impairments. The inclusion criteria were complete blindness or minimal light sensitivity with no ability to functionally use this sensation, as well as no pattern vision. Blind participants’ characteristics are presented in Table 1.

**Table 1.** Participant characteristics.

| Participant | Age<br>(years) | Sex | Cause of blindness | Age at blindness onset | Reading hand<br>(finger) |
| --- | --- | --- | --- | --- | --- |
| 1 | 24 | male | atrophy of the optic nerve | congenitally blind | left (index finger) |
| 2 | 26 | male | retinopathy of prematurity | congenitally blind | left |
| 3 | 37 | female | retinopathy of prematurity | congenitally blind | right |
| 4 | 28 | female | retinopathy of prematurity | congenitally blind | right |
| 5 | 25 | male | retinopathy of prematurity | congenitally blind | left |
| 6 | 34 | male | undefined (genetic) | congenitally blind | left |
| 7 | 32 | female | retinopathy of prematurity | congenitally blind | left |
| 8 | 43 | male | atrophy of the optic nerve | congenitally blind | left (index finger) |
| 9 | 31 | male | retinopathy of prematurity | congenitally blind | right |
| 10 | 32 | female | retinopathy of prematurity | congenitally blind | right (index finger) |
| 11 | 40 | female | atrophy of the optic nerve | congenitally blind | right (index finger) |
| 12 | 39 | female | retinopathy of prematurity | congenitally blind | left (index finger) |
| 13 | 40 | female | retinopathy of prematurity | congenitally blind | left |
| 14 | 30 | female | atrophy of the optic nerve | congenitally blind | right |
| 15 | 30 | male | optic nerve hypoplasia | congenitally blind | right |
| 16 | 39 | male | retinopathy of prematurity | congenitally blind | left |
| 17 | 27 | male | retinopathy of prematurity | congenitally blind | right |
| 18 | 45 | female | retinopathy of prematurity | congenitally blind | left |
| 19 | 45 | male | retinopathy of prematurity | congenitally blind | left |
| 20 | 22 | male | microphthalmia | congenitally blind | left |
| 21 | 45 | female | retinopathy of prematurity | congenitally blind | right (index finger) |
| 22 | 31 | female | atrophy of the optic nerve | congenitally blind | right (index finger) |
| 23 | 31 | male | retinopathy of prematurity | congenitally blind | left |
| 24 | 35 | female | congenital glaucoma | congenitally blind | both |
| 25 | 23 | male | atrophy of the optic nerve | congenitally blind | left (index finger) |
| 26 | 22 | male | retinopathy of prematurity | congenitally blind | left (index finger) |
| 27 | 33 | male | atrophy of the optic nerve | congenitally blind | left (index finger) |
| 28 | 29 | male | retinopathy of prematurity | congenitally blind | left (index finger) |
| 29 | 36 | female | undefined (genetic) | congenitally blind | right (index finger) |
| 30 | 42 | male | toxoplasmosis | congenitally blind | right (index finger) |
| 31 | 35 | female | undefined (genetic) | congenitally blind | right (index finger) |
| 32 | 40 | male | eye injury | 3 | right (index finger) |
| 33 | 23 | female | glaucoma | 4 | left (middle finger) |
| 34 | 26 | female | retinal detachment | 17 | left (index finger) |
| 35 | 38 | male | glaucoma | 21 | right (index finger) |
| 36 | 45 | female | eye injury | 23 | left (index finger) |

The study was approved by the Jagiellonian University Ethics Committee. All participants provided written informed consent before the study and were compensated for their time; blind participants’ travel expenses were reimbursed. The documents were read to blind participants by the experimenter, and the signature location was indicated with tactile markers.

### Experimental tasks and procedure

All volunteers were naïve to the experimental procedure. At the very beginning of the experiment and prior to the behavioral tasks, the participants were asked to fill out a questionnaire regarding their bodily experiences. Since increased physiological arousal has been shown to provide an advantage for heartbeat perception (Pollatos, Herbert, Kaufmann, Auer, & Schandry, 2007), to allow for any potentially elevated heart rates due to walking at a fast pace to the building, etc., to return to a normal level, we asked the participants to fill out the questionnaire at the beginning rather than the end of the procedure. The Multidimensional Assessment of Interoceptive Awareness (MAIA; Mehling et al., 2012; see Brytek-Matera & Kozieł, 2015 for a Polish translation and validation) is a 32-item tool that measures interoceptive body awareness, which consists of eight subscales, namely, *Noticing, Not-Distracting, Not-Worrying, Attention Regulation, Emotional Awareness, Self-Regulation, Body Listening*, and *Trusting*; the questionnaire has a range of scores of 0-5, with 0 indicating low and 5 indicating high interoceptive body awareness. For the same reason as that described above, i.e., to prevent potentially elevated heart rates, the participants were also asked not to consume any caffeinated drinks on the day of the experiment (see Hartley, Lovallo, & Whitsett, 2004; McMullen, Whitehouse, Shine, Whitton, & Towell, 2012).

Before the start of the behavioral part of the experiment, all the participants were informed about the experimental setup and received a short description of the procedure. Then, each participant sat on a chair in a comfortable position. A heart rate baseline reading was obtained over a period of 5 minutes before the beginning of the counting task. The participants’ heart rate was recorded using a Biopac MP150 BN-PPGED pulse oximeter (Goleta, CA, United States) attached to their left index finger and connected to a laptop with AcqKnowledge software (version 5.0), which recorded the number of heartbeats after the preset time. Then, the number of heartbeats was quantified using the embedded ‘count peaks’ function. To reduce the possibility that participants would perceive pulsation in their fingers due to the grip of the pulse oximeter, attention was given to ensure a comfortable and not overly tight fit of the finger cuff. Sighted subjects were blindfolded while performing the tasks.

Participants were given the following instructions: “Without manually checking, can you silently count each heartbeat you feel in your body from the time you hear ‘start’ to when you hear ‘stop’? Do not take your pulse or feel on your chest with your hand. You are only allowed to feel the sensation of your heart beating” (adapted from Garfinkel et al., 2015). After the trial, the participants verbally reported the number of heartbeats counted. They did not receive any feedback regarding their performance. Immediately after reporting the number of counted heartbeats, participants were asked to rate their confidence in the perceived accuracy of their response (see Garfinkel et al., 2015). This confidence judgment was made using a scale ranging from 0 (total guess/no heartbeat awareness) to 10 (complete confidence/full perception of heartbeat). A rest period of 30 seconds was given before the next trial began. The task was repeated six times to form six trials, using intervals of 25, 30, 35, 40, 45 and 50 seconds, presented in a random order. The participants received no information about the interval length.

To examine an additional dimension of metacognitive reflection, namely, prior and posterior beliefs of one’s performance (see Fleming, Massoni, Gajdos, & Vergnaud, 2016; Kirsch et al., 2021), after receiving the instruction of the task and being given an opportunity to ask clarifying questions, the participants were also asked to assess their prospective performance in the task in relation to all trials. Thus, before the task, they were given the following instruction: *Now that I explained you the task, how well are you going to perform in the task on a scale ranging from 0 (not so well/total guess) to 100 (very well/very accurate)?* After completing the task, participants were asked to reflect on their performance in all trials and were given the following instructions: *Now that you have done the task, how well did you perform in the task on a scale ranging from 0 (not so well/total guess) to 100 (very well/very accurate)?* These data were analyzed separately from the confidence judgments provided after every trial.

To examine a potential relationship between interoceptive and tactile abilities, we also employed a measure of tactile acuity, namely, the grating orientation task (Johnson & Phillips, 1981). The procedure followed the method described in Radziun et al. (2021; see also Supplementary Material). The grating orientation threshold was calculated by linear interpolation between grating widths spanning 75% correct responses (see Van Boven & Johnson, 1994; Merabet et al., 2008; Wong, Gnanakumaran, & Goldreich, 2011; Garfinkel et al. 2016b). Nine participants from the blind group and 12 participants from the sighted group were excluded from the data analysis because they could not perform the task beyond the expected level (75% accuracy).

After completing the tasks described in this study, the same participants also took part in two other behavioral experiments that examined body perception, which is not related to the current study’s research questions, and that will be reported separately (Radziun et al., in preparation).

### Data analysis

#### Interoceptive accuracy

For each participant, an accuracy score was derived, resulting in the following formula for interoceptive accuracy across all trials (see Schandry, 1981):

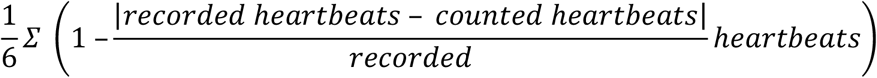

The interoceptive accuracy scores obtained following this transformation usually vary between 0 and 1, with higher scores indicating a better discrimination of the heartbeats (i.e., smaller differences between estimated and actual heartbeats). Two blind participants who failed to perform the task were excluded from further analyses (extremely low accuracy levels of –0.128 and –1.178; see also *Plan of statistical analysis*).

#### Interoceptive sensibility

MAIA scores served as an indication of the general interoceptive sensibility. Higher scores indicated higher interoceptive sensibility.

#### Interoceptive awareness

First, the mean confidence during the heartbeat detection task was calculated for every participant by averaging the confidence judgments over all the experimental trials to produce a global measure of mean confidence in perceived accuracy of response. Then, to provide an index of interoceptive awareness, a correlation coefficient between the accuracy score (see section *Interoceptive accuracy*) and the confidence ratings was calculated.

### Plan of statistical analysis

The data were tested for normality using the Shapiro–Wilk test and found to be not distributed normally (p < .05). Therefore, nonparametric statistics were used (Mann–Whitney *U* test for independent group comparisons and Spearman’s *rho* for correlations). All p values were two-tailed. Data exclusion criteria were established prior to data analysis.

For the Bayesian analyses, the default Cauchy prior was used. BF_01_ indicates support for the null over the alternative hypothesis, and BF_10_ indicates support for the alternative hypothesis over null hypothesis (e.g., a BF_01_ = 8 means 8 times more support for the null hypothesis, while BF_10_ = 8 means 8 times more support for the alternative hypothesis). BFs between 0.333 and 3 are normally considered inconclusive (Jarosz & Wiley, 2014; Lee & Wagenmakers, 2014).

The data were analyzed and visualized with RStudio software, version 1.3.1056, and the Bayes Factor software package, version 0.9.12-4.2.

### Data availability

All data generated and analyzed during the study are available from the corresponding author upon request.

## Results

### Interoceptive accuracy

Our results revealed that blind individuals had better interoceptive accuracy than sighted controls, as reflected by significantly higher performance in the heartbeat counting task (W = 836, p = 0.009, CI95% = 0.030–0.241, BF_10_ = 10.540; M_Blind_ = 0.779, SD_Blind_ = 0.166, M_Control_ = 0.630, SD_Control_ = 0.237; Figure 1). The baseline performance in the sighted control group was comparable to the results obtained in other studies using the heartbeat counting task paradigm (e.g., M = 0.66 in Garfinkel et al., 2015; M = 0.63 in Zamariola, Maurage, Luminet, & Corneille, 2018; M = 0.61 in von Mohr, Finotti, Villani, & Tsakiris, 2021), which highlights that the task was successfully implemented in the present study and that the blind group showed a level of accuracy that was significantly higher than the values normally reported in the literature.

**Figure 1.**
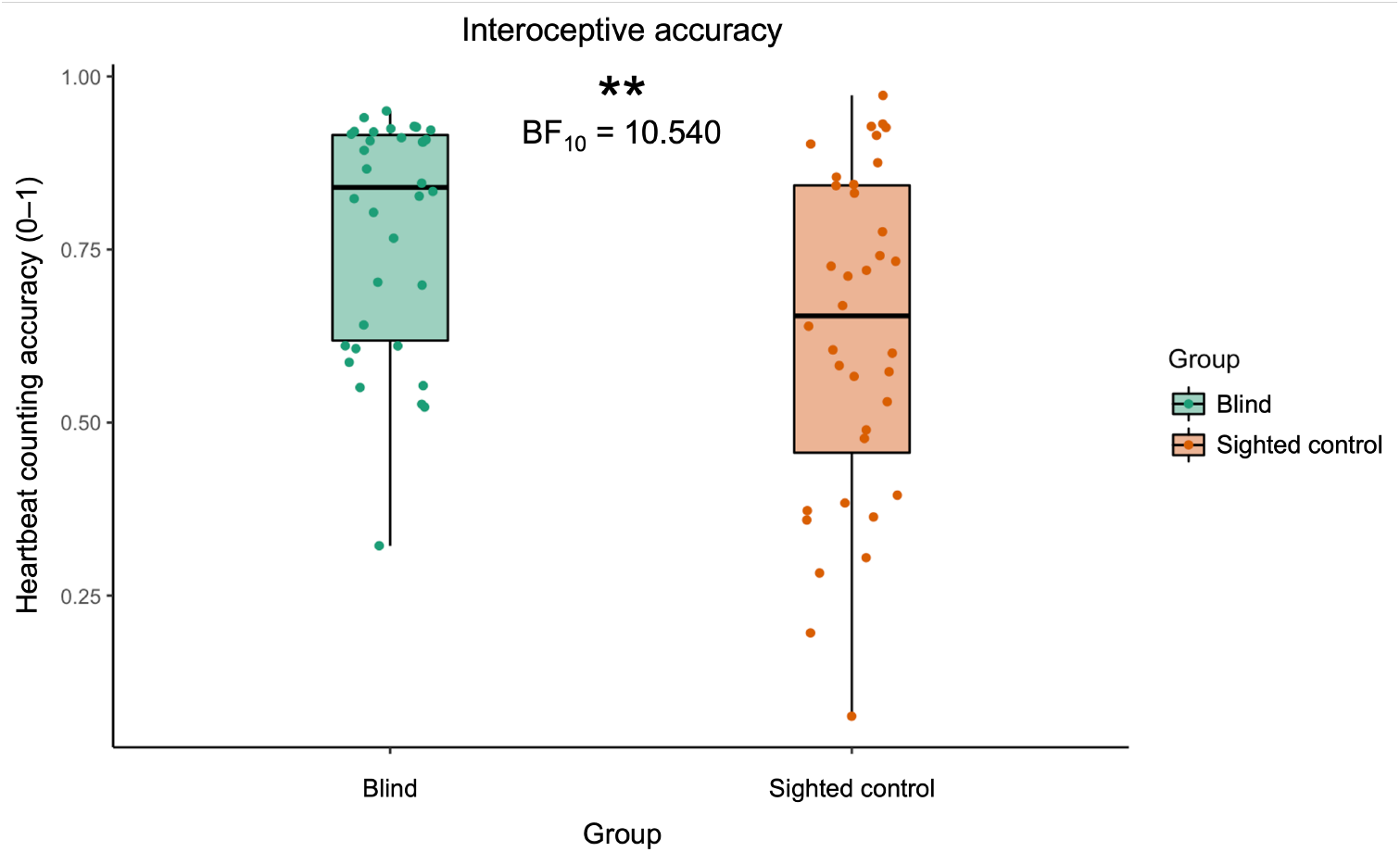
Interoceptive accuracy, as measured using the heartbeat counting task, was elevated in blind individuals compared to sighted controls. The boxplots depict the data based on their median (thick black line) and quartiles (upper and lower ends of boxes). The vertical lines, i.e., the whiskers, indicate the minimum or maximum values within 1.5x the interquartile range above and below the upper and lower quartiles. The datapoints outside the vertical lines are the outlier observations, the furthest being the minimum or maximum values in the data. The following figures are formatted in the same fashion.

The heart rate was equivalent for both groups (W = 569, p = 0.617, CI95% = −6.000– 3.400, BF_01_ = 3.479; M_Blind_ = 76.347, SD_Blind_ = 10.441, M_Control_ = 77.794, SD_Control_ = 9.663). Therefore, the potential influence of heart rate, which has been shown to be a factor that affects performance (e.g., Radziun et al., 2021), could be excluded as an explanation for the effect observed here.

### Interoceptive sensibility

There was no significant difference in average MAIA scores between the groups (W = 607, p = 0.958, CI95% = −0.283–0.323, BF_01_ = 4.046; M_Blind_ = 2.885, SD_Blind_ = 0.643, M_control_ = 2.900, SD_Control_ = 0.561; Figure 2), which shows that there was no difference in subjective interoceptive sensibility between the blind group and the sighted control group. No significant differences between the groups emerged when comparing the MAIA subscales separately (all p > 0.05).

**Figure 2.**
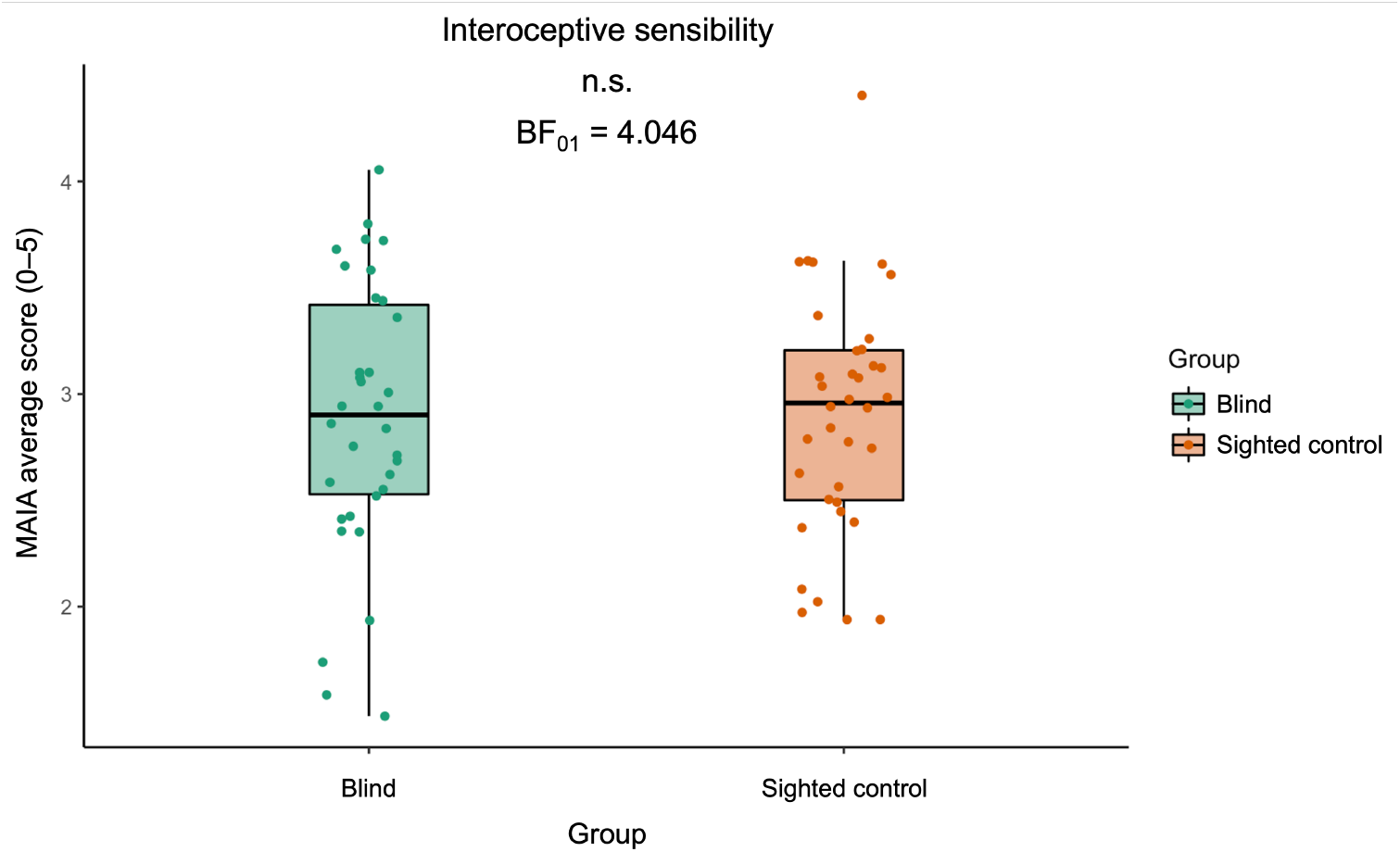
There was no difference between the blind group and the sighted control group in interoceptive sensibility, as measured with the Multidimensional Assessment of Interoceptive Awareness (MAIA).

Interoceptive sensibility, as measured by the MAIA, and interoceptive accuracy did not correlate in either the blind group (ϱ = 0.183, p = 0.298, CI95% = −0.165–0.491, BF_01_ = 2.223) or the sighted controls (ϱ = 0.253, p = 0.136, CI95% = −0.082–0.537, BF_01_ = 1.517), which suggests that subjectively reported sensitivity to bodily sensations does not align with objective interoceptive accuracy regardless of the visual experience; this follows a pattern that has been observed in previous studies (e.g., Mai, Wong, Georgiou, & Pollatos, 2018).

### Interoceptive awareness

In the blind group, we did not find a significant correlation between interoceptive accuracy, as measured by heartbeat counting task, and interoceptive sensibility, as measured by the average confidence ratings (ϱ = 0.277, p = 0.113, CI95% = −0.067–0.563, BF_01_ = 0.362; Figure 3A). However, this correlation was found in the sighted control group (ϱ = 0.484, p = 0.003, CI95% = 0.185–0.701, BF_10_ = 39.449; Figure 3B).

**Figure 3.**
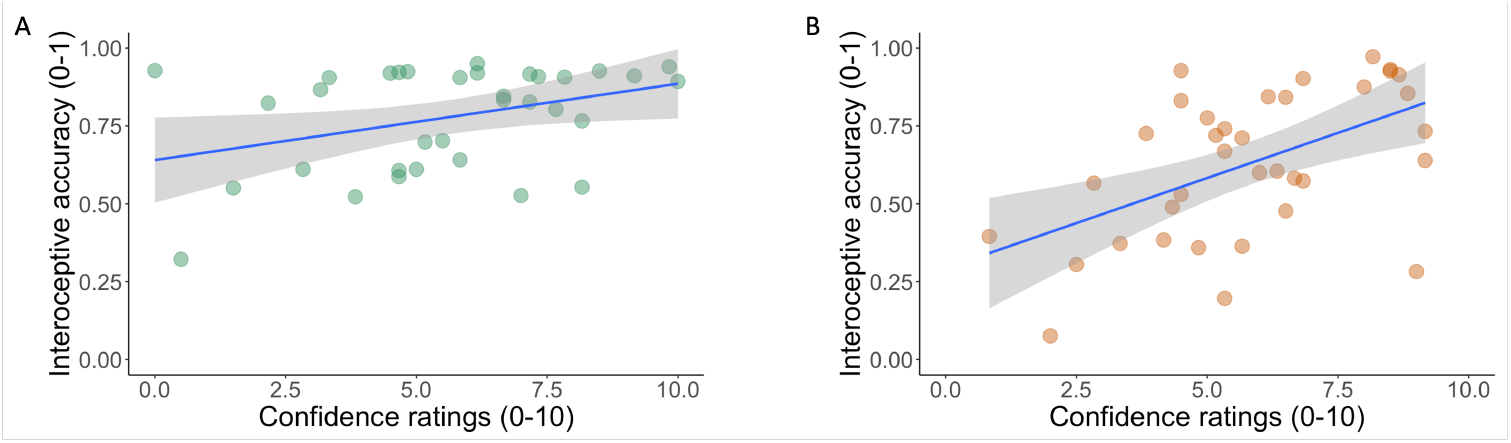
Confidence–accuracy correlation in the blind group (A) and the sighted control (B) group.

Notably, there was no significant difference in the mean confidence ratings between the blind group and the sighted control group (W = 601.5, p = 0.906, CI95% = −1.333–1.000, BF_01_ = 3.881; M_Blind_ = 5.637, SD_Blind_ = 2.501, M_Control_ = 5.819, SD_Control_ = 2.145; Figure 4).

**Figure 4.**
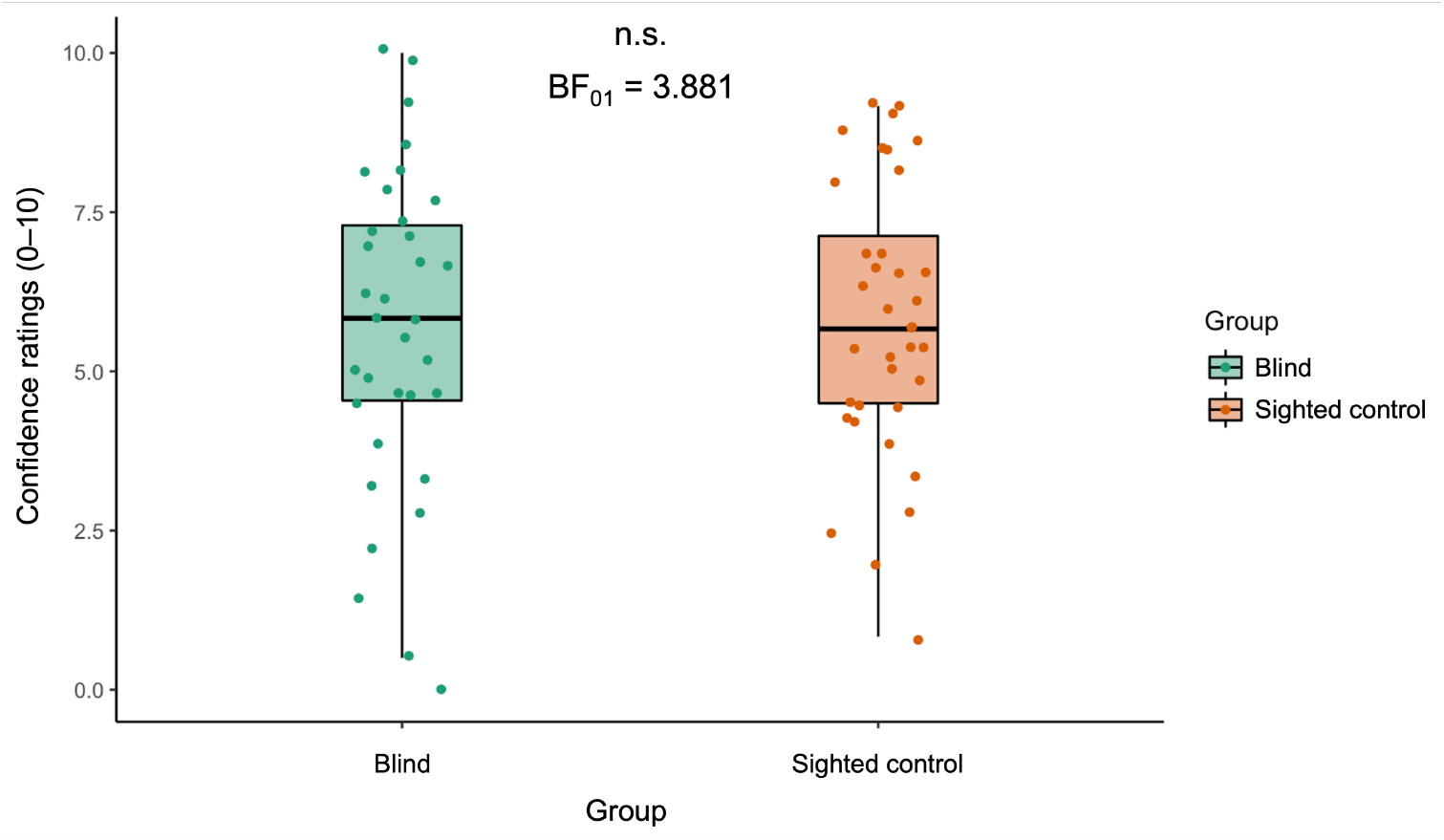
There was no difference between the blind and sighted control groups in the average confidence ratings.

### Belief of performance accuracy

We found no difference between the blind and sighted control groups in regard to their belief in the performance of accuracy, for both completion before the task (W = 460.5, p = 0.075, BF_01_ = 1.129; M_Blind_ = 50.588, SD_Blind_ = 27.900, M_Control_ = 61.472, SD_Control_ = 24.455) and completion after the task (W = 543, p = 0.419, BF_01_ = 3.006; M_Blind_ = 51, SD_Blind_ = 26.212, M_Control_ = 56.083, SD_Control_ = 24.450).

### Relationship between interoceptive accuracy and tactile acuity

We found no correlation between interoceptive accuracy and tactile acuity in either the blind (ϱ = −0.209, p = 0.293, CI95% = −0.546–0.185, BF_01_ = 1.954) or sighted control (ϱ = - 0.101, p = 0.640, CI95% = −0.484–0.316, BF_01_ = 2.047) groups (Figure 5).

**Figure 5.**
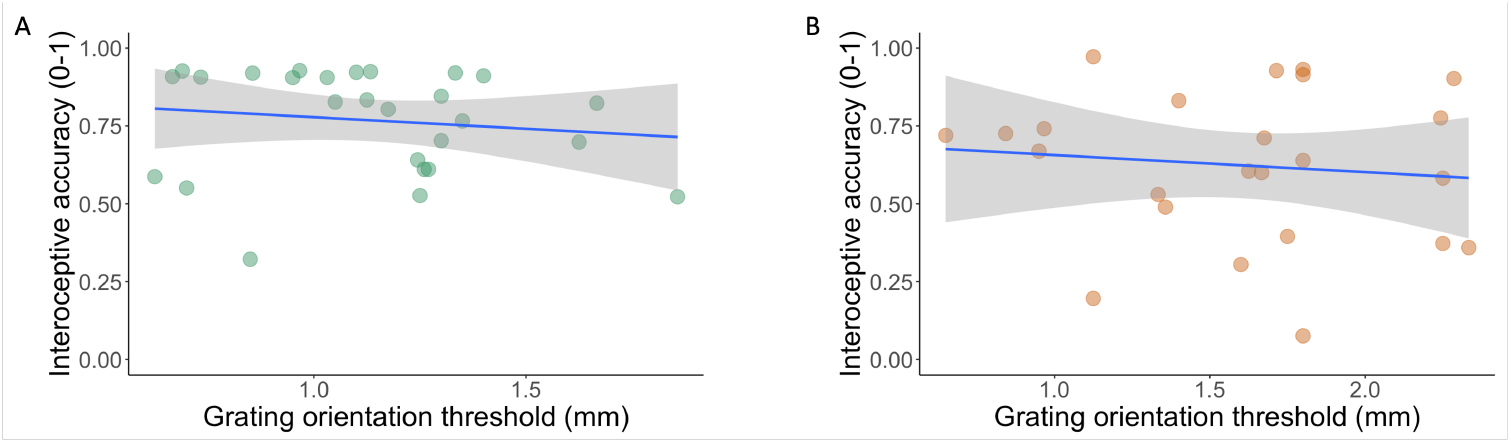
Correlation between interoceptive accuracy and tactile acuity in the blind (A) and sighted control (B) groups.

## Discussion

In this study, we investigated the effect of blindness on cardiac interoception. Consistent with our hypothesis, we found that the blind group performed better than the sighted control group on the heartbeat counting task; that is, that the blind individuals had better cardiac interoceptive precision compared to the control group. Interestingly, this effect appears to pertain only to pure sensory abilities; we did not find any differences in regard to interoceptive sensibility as measured by a subjective questionnaire, namely, the MAIA. We also did not find differences in confidence in the given response or belief of performance accuracy, which were measured both before and after task completion. We did not find any differences in heart rate either, which excludes the possibility that the observed effect was due to a potential discrepancy between the groups occurring at the physiological level. Taken together, our results suggest that blind people are better able to sense their own heartbeats compared to their sighted counterparts.

The reasons behind our main result could be twofold. On the one hand, this result could reflect a genuinely increased perceptual ability to use the visceral information from rhythmic cardiovascular events felt in the chest, which leads to a more accurate counting of heartbeats. This is the most straightforward and the most likely interpretation, especially considering the results of the tactile control task. An alternative interpretation that we cannot exclude is that blind individuals showed a more accurate performance in the task because they were better at sensing pulsations from different locations in their body (see also Betka et al., 2021) and picking up subtle cues from forehead, limbs, etc., thus relying more on multisensory integration of various somatosensory and interoceptive signals related to the heartbeats rather than sensory signals from the heart that are mediated through the vagal nerve (Prescott & Liberles, 2022). Future studies should investigate this further; however, in either case and regardless of the underlying sensory mechanism, the current results are important because they suggest that in general, blind individuals are better at perceiving their heartbeats than sighted individuals.

What kind of mechanism could trigger the kind of cross-modal plasticity that would lead to improvements in cardiac interoception? Several studies with blind individuals have suggested that their improved sensory acuity is not necessarily driven by the lack of vision itself, but rather due to the experience-dependent neuroplastic mechanisms—caused by, for example, the increased training of the hands due to tactile exploration of everyday objects and Braille reading (Alary et al., 2009; Sathian & Stilla, 2010, Voss, 2011; Wong et al., 2011). However, such an explanation seems unlikely for the enhancements in cardiac interoceptive accuracy observed in our study. Tactile training among blind individuals is predominantly involuntary and associated with exploring the environment and performing various daily activities, while interoceptive functions are usually not trained in this way. A potential interoceptive equivalent of tactile training could be the practice of meditation. However, previous research has suggested that regular meditation does not lead to superior interoceptive accuracy (e.g., Khalsa et al., 2008; Khalsa, Rudrauf, Hassanpour, Davidson, & Tranel, 2020; see also Farb, Segal, & Anderson, 2013). Given that the experience-dependent explanation of the effect observed in our study seems unlikely, the results fit better in the theoretical framework of massive cross-modal plasticity occurring because of visual deprivation itself. In this view, the lack of visual experience leads to neuroplastic changes in sensory, multisensory, and visual areas and their anatomical interconnections that provide greater neural processing capacities for the remaining senses, including cardiac interoception, as has been revealed by the current results. The fact that such massive cross-modal plasticity effects go beyond the exteroceptive senses of hearing, discriminative touch, and olfaction to include sensations from an inner visceral organ is particularly noteworthy, as it advances our understanding of the extent of such effects and related perceptual enhancements.

What could be the neuroanatomical basis for the current findings of enhanced cardiac interoceptive accuracy? One of the regions that is important for the processing of afferent visceral information, including cardiac signals, is the anterior insula (see Critchley et al., 2004). Interestingly, visual deprivation has recently been found to reshape the functional architecture within anterior insular subregions (Liu et al., 2017). Although it is not clear how these neuroplastic changes are related to the ability to perceive heartbeats or other visceral signals, future neuroimaging studies could explore this possible link. Furthermore, the observed enhancement could also be due to structural changes within the deprived occipital cortex. Indeed, previous studies have reported a relationship between increased occipital cortical thickness and enhanced performance within the auditory modality (Voss & Zatorre, 2012). Future studies might elucidate the relationship between structural changes in the brains of blind individuals and their superior performance in sensory tasks.

Future studies should also investigate to what extent the current findings are generalizable to other interoceptive submodalites, such as the processing of information from other inner organs (e.g., bladder and lungs) and including skin-based interoception (see Crucianelli & Ehrsson, 2022). Interestingly, pain detection thresholds have been found to be lower in blind individuals, which indicates that they are more sensitive to detecting nociceptive stimuli on the skin (Slimani et al., 2013; Slimani et al., 2014; Slimani, Ptito, & Kupers, 2015; Slimani, Plaghki, Ptito, & Kupers, 2016), thereby paralleling findings in animal models of blindness (Touj, Tokunaga, Al Aïn, Bronchti, & Piché, 2019; Touj, Paquette, Bronchti & Piché, 2021). It has also been proposed that blind individuals might rely more on internal hunger cues rather than taste in food choices (Gagnon, Kupers, & Ptito, 2013). Although it may suggest that the current findings could be generalizable, recent studies in sighted individuals have found that perceptual abilities on different interoceptive tasks that probe different submodalites are independent and uncorrelated (Garfinkel et al., 2016b; Ferentzi et al., 2018; Crucianelli, Enmalm, & Ehrsson, 2021) and that cardiac interoceptive accuracy does not correlate significantly with accuracy measures in skin-based interoceptive tasks (Crucianelli et al., 2021). Thus, to obtain a more complete picture of how visual deprivation affects interception, future studies should employ a battery of tests and investigate how different interoceptive submodalities are affected by visual deprivation and if some are more influenced than others.

Surprisingly, in our study, we did not observe a significant correlation between interoceptive accuracy and interoceptive sensibility (as measured by confidence ratings) in blind individuals, although the Bayesian analysis suggested that this result could be inconclusive. In the sighted group, in turn, this correlation was positive, significant and supported by Bayesian statistics. In previous studies, higher levels of interoceptive accuracy have been associated with higher interoceptive awareness and lower interoceptive accuracy with no relationship between accuracy and sensibility (e.g., Garfinkel et al., 2015; García-Cordero et al., 2016; Murphy, Catmur, & Bird, 2018). In other words, healthy sighted people who do well on the heartbeat counting task also have a metacognitive awareness that they are doing well, whereas individuals who perform poorly also do less well in judging how poor their performance is. The present findings may indicate that this relationship is different in blind individuals, which suggests a lowered insight into sensory abilities. This would also be consistent with the results from the MAIA questionnaire that suggested no differences in how the blind and sighted participants rated a range of sensations related to various aspects of interoception in their daily life. However, Beaulieu-Lefebvre and colleagues (2011) reported that blind individuals scored higher than sighted individuals on a scale that assessed sensibility to olfactory sensations, although subsequent studies did not find conclusive evidence for the difference between blind and sighted individuals in metacognitive abilities in relation to olfactory task performance (Cornell Kärnekull, Arshamian, Nilsson, & Larsson, 2016). Future studies should try to clarify whether insight into perceptual abilities among blind people varies between interoceptive and exteroceptive senses.

It is well known that internal bodily signals—cardiac signals in particular—are in a mutual interactive relationship with emotion processing (see Pollatos, Gramann, & Schandry, 2007; Critchley & Harrison, 2013; Adolfi et al., 2016; Garfinkel & Critchley, 2016c; Critchley & Garfinkel, 2017; Shah, Catmur, & Bird, 2017). Changes in afferent interoceptive inputs from the heart modulate subjective emotions (e.g., the intensity of experience fear; see Garfinkel et al., 2014), and changes in emotion can trigger various physiological peripheral reactions in the body (e.g., increasing heart rate), which in turn modulate the ascending interoceptive signals in the brain. Thus, enhanced cardiac interoception in blind individuals may modulate these body-brain interactions and lead to changes in emotional processing. Furthermore, it has been suggested that the degree to which an individual is able to recognize their own interoceptive states positively correlates with how well they recognize emotions in themselves and in others (Wiens, Mezzacappa, & Katkin, 2000; Herbert, Pollatos, & Schandry, 2007; Terasawa, Moriguchi, Tochizawa, & Umeda, 2014; Ernst et al., 2014; Shah, Hall, Catmur, & Bird, 2016; Bird, Shah, & Catmur, 2017; Tsakiris, 2017; but see also Ainley, Maister, & Tsakiris, 2015). Blind individuals do not show impairments in emotion processing (Gamond, Vecchi, Ferrari, Merabet, & Cattaneo, 2017); moreover, they show better discrimination of emotional information, along with increased amygdala activation to emotional auditory stimuli (Klinge, Röder, & Büchel, 2010; Klinge, Röder, & Büchel, 2013), where the amygdala, along with the insula, is one of the critical structures for interoception and emotion processing (Critchley, Mathias, & Dolan, 2002). Therefore, our results could provide a missing explanatory link between improved emotional processing and increased sensory acuity in the blind.

Our results could also have important implications for future research on bodily awareness and self-consciousness in the blind. Heartbeats are one of the first sensory cues during early development, occurring after 5 ½ to 6 weeks after gestation; even young infants seem to perceive their own heartbeats, as demonstrated in behavioral paradigms (Maister, Tang, & Tsakiris, 2017; but see also Weijs, Daum, & Lenggenhager, 2022). Thus, together with proprioceptive feedback from movements, cardiac interoceptive signals may play an important role in the developing central nervous system in regard to laying the foundation for sensory processing and the sense of self (see also Quigley, Kanoski, Grill, Feldman Barrett, & Tsakiris, 2021). In sighted individuals, visual experience later becomes crucial when the infant learns to interact with external objects and recognizes their own body parts through movement and visuotactile feedback (Rochat & Striano, 2000; Zmyj, Jank, Schütz-Bosbach, & Daum, 2011; Chen, Lewis, Shore, Spence, & Maurer, 2018). These visual experiences of the self and the world presumably drive the development of a multisensory sense of the bodily self (Bremner, 2016); however, in blind individuals who lack this kind of information, interoception may play a relatively greater role. It has been shown that congenitally blind individuals exhibit changes in the multisensory representation of their own body (Petkova, Zetterberg, & Ehrsson, 2012; Nava, Steiger, & Röder, 2014). Thus, the current findings might be important for future research into how bodily awareness and self-consciousness develop and are maintained without vision and how enhanced ability to sense cardiac signals may modulate bodily awareness and self-consciousness in the blind.

In conclusion, we have conducted the first study on cardiac interoceptive abilities in blind individuals and found that blind individuals are better than their sighted counterparts at sensing their own heartbeats in the classic heartbeat counting task. The results can contribute to our understanding of the fundamental constraints of massive cross-modal plasticity after blindness by suggesting that visual deprivation leads to interoceptive plasticity, which may have interesting potential implications for emotional processing, bodily awareness, and the conscious experience of the self.

## Acknowledgments

We thank Adam Enmalm for his assistance in data collection during the pilot stage of the study.

