## Supplementary Material for "Cardiac interoception is enhanced in blind individuals"

Radziun et al.

**Supplementary Information.**

Grating orientation task

The stimuli used in this procedure were composed of eight hemispheric plastic domes that were stamped with equally wide parallel bars and grooves (JVP [Johnson-Van Boven-Phillips] Spatial Discrimination Domes, Stoelting, Inc. Wood Dale, IL), with widths of the following sizes: 0.35, 0.5, 0.75, 1, 1.2, 1.5, 2 and 3 mm. During the task, the right index finger of the participant was fixated on a table in a palm-up position. Gratings were applied with moderate force by a trained experimenter to the distal pad of the right index finger for ~1.5 s. The experimenter took care to avoid any movement of the participant’s finger caused by contact with the grating. The stimuli were applied in either horizontal or vertical manner relative to the long axis of the finger. A two-alternative forced-choice paradigm was used in which participants were asked to report whether the orientation of the grating was horizontal or vertical. The task consisted of eight blocks, with one for each grating width, while each block consisted of 20 randomized trials, half with gratings presented horizontally and half with gratings presented vertically. The order of blocks was fixed and corresponded to decreasing width of the gratings. No feedback about the accuracy of the response was given to the participants at any time.
